# Visual crowding is a combination of an increase of positional uncertainty, source confusion, and featural averaging

**DOI:** 10.1101/088898

**Authors:** William J Harrison, Peter J Bex

## Abstract

Although we perceive a richly detailed visual world, our ability to identify 1 individual objects is severely limited in clutter, particularly in peripheral vision. Models of such crowding have generally been driven by the phenomenological misidentifications of crowded targets: using stimuli that do not easily combine to form a unique symbol (e.g. letters or objects), observers typically confuse the source of objects and report either the target or a distractor, but when continuous features are used (e.g. orientated gratings or line positions) observers report a feature somewhere between the target and distractor. To reconcile these accounts, we develop a hybrid method of adjustment that allows detailed analysis of these multiple error categories. Observers reported the orientation of a target, under several distractor conditions, by adjusting an identical foveal target. We apply new modelling to quantify whether perceptual reports show evidence of positional uncertainty, source confusion, and featural averaging on a trial-by-trial basis. Our results show that observers make a large proportion of source-confusion errors. However, our study also reveals the distribution of perceptual reports that underlie performance in this crowding task more generally: aggregate errors cannot be neatly labelled because they are heterogeneous and their structure depends on target-distractor distance.

---

Throughout the entire visual field, vision is constrained by multiple bottlenecks in visual processing that limit the information reaching our awareness. Initially, information is lost to physiological factors such as the eyes' optics and retinal nerve fiber density, and neural selective sensitivity to spatio-temporal patterns ^1,2^. However, our ability to identify even a simple object, such as a letter or an oriented grating, is far worse than predicted from these factors when the object is surrounded by clutter ^3,4^. These identification failures, referred to as “crowding”, occur even though adaptation after-effects demonstrate that the object's features have been encoded, at least in primary visual cortex ^5-7^. Thus, our ability to consciously access the identity of an object is constrained by information processing capacity, not simply by retinal physiology or sensitivity limitations of the visual system. ^28^

Crowding, the inability to recognise an object in visual clutter, influences many aspects of vision. It is generally agreed to occur across the entire visual field ^8^, although it is markedly more difficult to measure at the fovea ^9^. As discussed in a review by Pelli and Tillman ^4^, crowding affects all basic object recognition tasks, predicts reading speed and dyslexia, and is diagnostic of foveal deficits present in amblyopia ^10^. Furthermore, it limits visibility of naturalistic images ^11-13^, and interacts with saccadic and smooth pursuit eye movements in non-trivial ways ^14-19^. There have been several recent reviews of crowding that summarize very well its ubiquity ^4,20-22^, as well as examples in which object recognition is seemingly unaffected by parameters that cause crowding in other instances e.g. ^23^.

The spatial extent of crowding is quite similar across paradigms. Perceptual errors increase with eccentricity and decrease as the distance between target and distractor increases. The precise target-flanker distance at which crowding is alleviated at a given eccentricity, often referred to as Bouma's constant, is somewhat variable across studies ^8^, and changes dynamically according to the duration and relative timing of target and distractors ^19,24-28^. Despite this variability, the term “crowding” is typically taken to refer to any target identification interference that depends on target-distractor proximity ^29^.

Although there are many fascinating aspects of crowding, we focus on fundamental findings. By viewing Figure 1, the reader can experience firsthand three phenomenal aspects of crowding that arise with stimuli similar to those used in our experiments. Following standard convention, we term these phenomena: 1) positional uncertainty ^30,31^, 2) feature averaging ^32^, and 3) source confusion ^33,34^. In this figure we present the same target stimulus, a Landolt C ^1^, in a series of distractor conditions. An observer's goal is to locate the orientation of the gap section as accurately as possible. If the reader fixates the spots in succession from top to bottom, they may note that the apparent clarity of the correspondingly-coloured target orientation is affected differently in each condition. The target gap is clearest in the top row, but its position is less clear when fixating the yellow spot below, perhaps because the solid ring distractor adds noise to the positional mechanisms encoding the target orientation ^35^. When fixating the pink spot, the target and distractor gaps may perceptually blend together, shifting the perceived target orientation toward the distractor orientation e.g. ^36^. When fixating the green spot, it is not immediately clear which of the multiple gaps is the target, and it may be easy to confuse a distractor gap for a target gap ^37^. The changes in target visibility while viewing the yellow, pink, and green stimuli demonstrate, in order, positional uncertainty, feature averaging, and source confusion.

**Figure 1.**
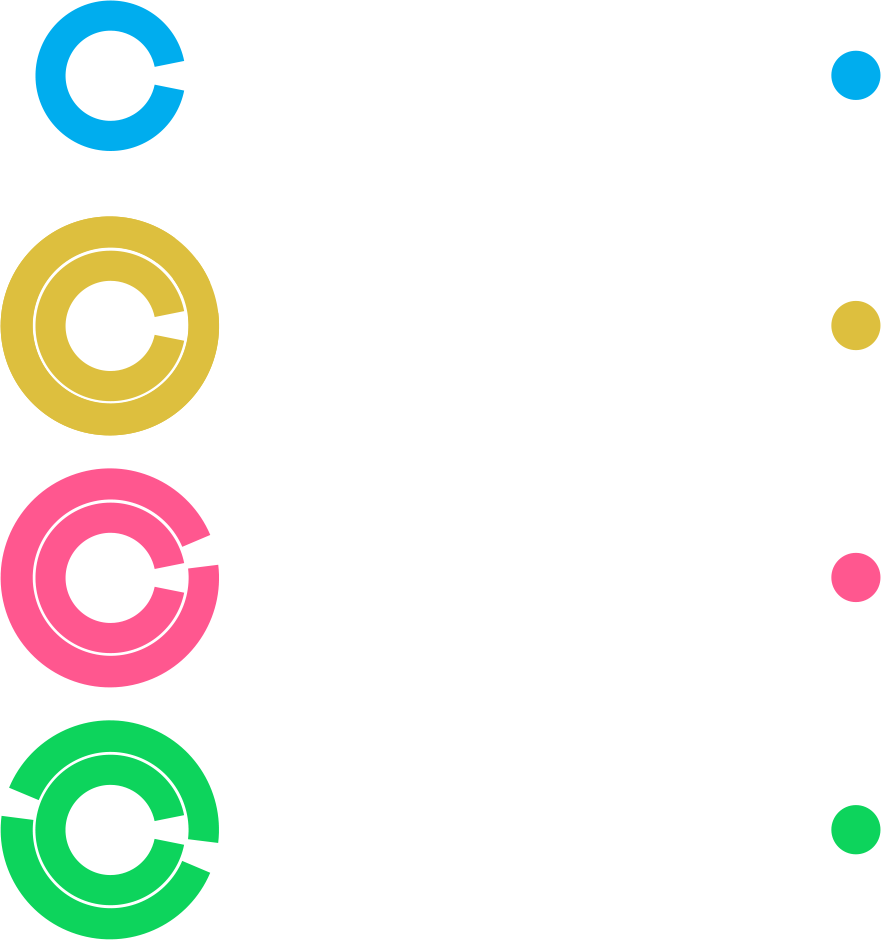
Examples of stimuli in our experiments that produce qualitatively different perceptual outcomes. From top to bottom, rows represent the unflanked, no-gap flanker, one-gap flanker, and two-gap flanker conditions. Stimuli are drawn to scale, but were white on a grey background in the experiment. Data in later figures follow the colour conventions shown here.

The perceptual phenomena experienced in cluttered displays are typically revealed across studies employing different methodologies. Changes in positional uncertainty and feature averaging are found in experiments in which an observer is required to make a spatial judgment about a continuous property of the target, such as its orientation or relative position, for examples, see ^36,38^.

Feature substitutions – mistaking a distractor element for a target – are mostly found in paradigms in which the observer is required to report the categorical identity of target such as a letter; trials in which the observer reports a distractor identity instead of the target reveal source confusions ^33,34,39^, which may be independent of an increase in positional uncertainty ^40-42^.

It is important to note that distinct categories of errors, such as *averaging* and *substitution* errors, are descriptors of results, not descriptors of a mechanism per se. Indeed, even the term crowding refers to the result of some visual process and not a mechanism. The underlying cause of crowding has previously been explained by various computational models ^11,22,36-38,43-45^ and higher-level mechanistic hypotheses ^23,46^. Population code models, in which all visual features probabilistically contribute to perceptual reports, can produce a wide variety of data ^47^, including so-called averaging and substitution errors ^37^. We have thus argued that the different classes of errors reported across the crowding literature are actually arbitrary categories of the output of a single mechanism. In the present report, therefore, we use the terms “substitution” and “averaging” as a convenient way to describe patterns in our data, but not to indicate hypothesised mechanisms. Our aim in the present study is to shed further light on the cause of crowding using a single paradigm that produces multiple perceptual phenomena. Here we use experiment and modelling to quantify changes in positional uncertainty, averaging, and source confusions in visual clutter.

## Methods

This experiment accorded with the protocols reviewed and approved by our local institutional review board. We tested three highly experienced psychophysical observers, including the two authors, all of whom gave informed consent. In figures, we refer to the participant naïve to the specific purposes of this experiment as N1, and to the authors as A1 (PJB) and A2 (WJH). All observers previously participated in two similar crowding experiments ^37^.

Observer sat 57 cm from the display with their head stabilised by a chin and headrest. The display was a CRT monitor (1280 × 1024 resolution, 85 Hz). We programmed the experiment with the Psychophysics Toolbox Version 3 ^48,49^ in MATLAB (MathWorks). Stimuli were white (100 cd/m^2^) on a gray (50 cd/m^2^) background. The target was centered 10° to the right of the fixation spot, had a 2° diameter, and a line width of 0.4°. The gap width, measured at the midpoint of the line width, was 0.4°. In the one-gap and two-gap flanking conditions, the size of the gaps remained constant across flanker diameters. For all flanking conditions, the line width remained constant (0.4°). A flankers' outer edge was separated from the target's outer edge by 0.4°, 0.92°, 1.62°, 2.58°, 3.9°, or infinity (ie. no flanker). We express flanker size as the flanker radius as a proportion of the eccentricity of its centre, giving 0.14 ϕ, 0.19 ϕ, 0.26 ϕ, 0.36 ϕ and 0.49 ϕ, where ϕ is the flanker eccentricity (10°). The flanker condition was selected randomly from trial-to-trial.

The orientations of flankers were constrained in the following way to produce maximal crowding effects ^37^. In the one-gap and dual-gap flanker conditions, the flanker orientation was drawn from a normal distribution, centred on the target orientation and with a standard deviation of 22.5°. Within this range, we expect maximum levels of crowding. For the dual-gap flanker condition, a second flanker gap was drawn from a normal distribution centred 180° from the first flanker gap, with a standard deviation of 22.5°.

Each trial began when an observer pressed the space bar, which triggered the display of a small spot in the centre of the screen and the target (with or without flanks) for 500 ms. Immediately following the offset of the target, a Landolt C was presented in the centre of the display, and observers could rotate this clockwise or anti-clockwise by pressing the right or left arrow key respectively. To report an orientation, they pressed the space bar, and the next trial would begin. Observers were instructed to be as accurate as possible without rushing. They could take a break at any time by withholding a report. All observers completed 320 trials per session (20 repetitions of each target-flanker combination), for five sessions, giving a total of 1600 trials, or 100 trials per target-flanker condition. Each session took approximately 15 – 20 minutes.

## Results

For all conditions, report error is defined as the difference between the reported orientation and the target orientation, with positive errors indicating a report that was more clockwise than the target. Because in our experiments increasing the flanker size increases target-flanker separation, we use the terms “flanker size” and “target-flanker separation” interchangeably.

### No-gap flanker condition

Because the no-gap flanker has no features that could be substituted or averaged with the target, results from this condition allow us to examine clearly changes in positional uncertainty. In the no-gap flanker condition, observers' report errors clustered around 0° for all target-flanker separations (Fig. 2A). We used the Circular Statistics Toolbox^2^ to find the circular standard deviation of observers' reports. We refer to this measure as perceptual error, which is plotted in degrees separately for each observer in Figure 2B-D as a function of the flanker size. Consistent with the crowding literature, all participants' perceptual error was higher than the unflanked condition (dashed lines) for the two smallest flanker sizes. Observers N1 and A2 in particular show the characteristic, approximately linear improvement in performance with increasing flanker size.

Performance in the largest flanker condition was similar to unflanked performance for all observers.

**Figure 2.**
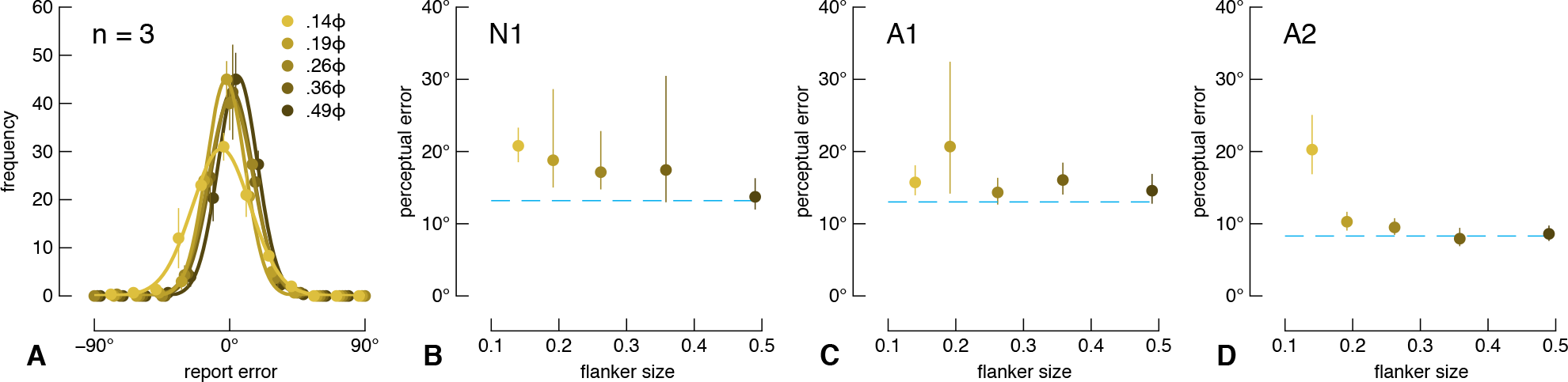
Results from the no-gap flanker condition. A) Average frequencies of each report error size (25 bins; bin width = 15°), fit with circular distributions. To improve visibility, the data have been offset slightly along the x-axis, and the x-axis has been truncated to ±90°. Only a total of 4 errors greater than ±90° were made by all observers across all no-gap flanker conditions. The inset legend shows the target-flanker separation condition. Error bars show one standard error. B – D) Perceptual error (the circular standard deviation of report errors) plotted as a function of flanker size for each participant. Flanker size is expressed as flanker radius divided by the eccentricity of its centre (i.e. in units of Bouma's constant). The dashed line indicates perceptual error for the unflanked condition. Error bars show 95% bootstrapped confidence intervals.

### One-gap flanker condition

In the one-gap flanker condition, we have previously shown that observers' reports correspond to the distribution of weighted responses to the target and flanker orientations ^37^. Thus, reports may include a proportion of responses at the target orientation, the flanker orientation and their mean orientation. Quantifying reports with a unimodal circular standard deviation measure as used for the no-gap flanker condition above, is inappropriate in such a case. For a detailed explanation from the working memory literature, see ^50^. We express report errors as a function of target-flanker orientation difference (see Fig. 3A and Appendix Figure A1A). Under a first analysis (see Appendix A), we fit linear models to the data: reports at the target orientation have a slope of zero, reports at the flanker orientation fall on the diagonal with unity slope, and reports at the average orientation fall on the diagonal with a slope of 0.5. For all observers for the two most crowded conditions, the slopes were close to 0.5 and reduce to 0 at larger target-flanker separations. However, if these data were composed of noisy target and noisy flanker reports as described above and as has been argued previously ^41^, the slope of a linear fit to crowded data may give a spurious interpretation favoring the averaging model. We further applied maximum likelihood mixture modelling as described by Bays et al ^50^, as well as Monte Carlo simulations, but these alternative analyses failed to return the true proportions of underlying report types of simulated data with known distributions. We thus used a simplified approach that labels each datum according to its distance from each model prediction, as described below.

To quantify report errors in the one-gap flanker condition, we measured the distance of all report errors from each of three underlying model predictions that correspond to target reports, averaged reports, or substitution reports, and labelled each datum according to the nearest model (see Fig. 3A). We then quantified report types as a proportion of all trials from each condition. These proportions are shown for all target-flanker separations in Figure 3B-D with symbols indicating observers as per the legend. The ordinate labels on the different panels correspond to different model predictions. Note that this analysis fits data simultaneously to all model predictions, and so summing across panels for one flanker size for a single observer gives 1. The pattern is very similar for all observers: with increasing flanker size, the proportion of target reports increases, the proportion of average reports is relatively stable, and the proportion of substitution reports decreases. Note that this pattern of results indicates that the response distribution is multi-modal, since the proportions of data for each error type changes non-monotonically. Although the proportion of target reports saturates around 0.6 (Fig. 3B), this is likely an underestimate due to a limitation of our analysis for distributions when the response standard deviation is large and the target-flanker orientation difference is small: our simulations revealed that the modelled proportion of target reports is accurate when the proportion of other model components is high (greater than ~0.3), but the proportion of target reports is underestimated when the contribution from other model components is minimal. For small flanker sizes, the conditions under which we expect relatively poor performance, the proportion of each report type is likely more accurate than for larger flanker sizes. Based on our previous work and the crowding literature, it is likely that observers' reports are barely, if at all, influenced by the flanker for the largest flanker condition (for example, see Fig. 2).

**Figure 3.**
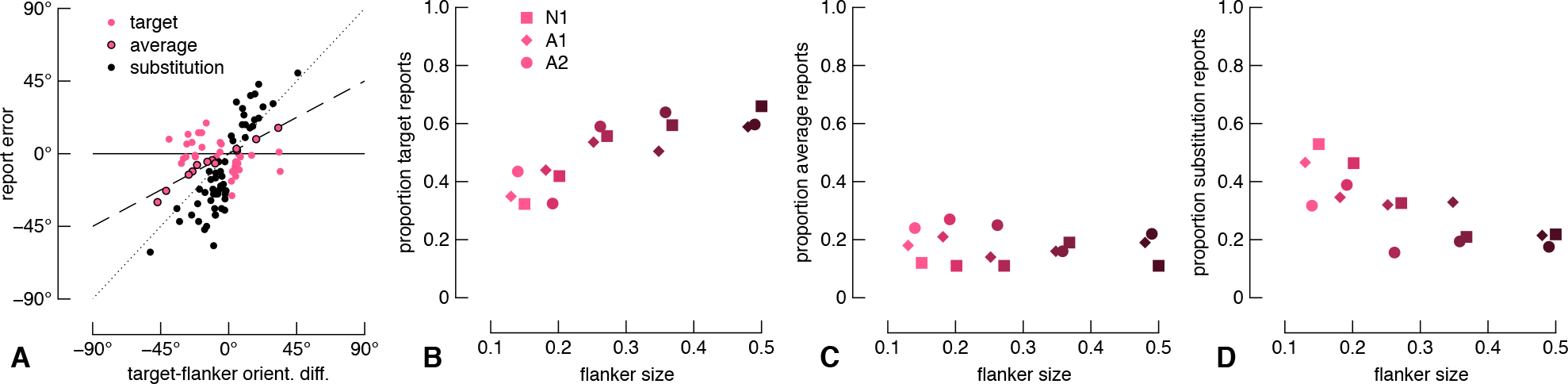
Results from the one-gap flanker condition. A) Raw report errors from the naïve participant. Solid, dashed, and dotted lines show target, averaging, and substitution models, respectively. Data are attributed to each model according to proximity. B-D) Proportion of report type as a function of flanker size for all observers. Report type is shown on the ordinate across panels. Data for different observers have been horizontally offset slightly for clarity.

### Two-gap flanker condition

In the two-gap flanker condition, one flanker gap orientation was normally distributed around the target orientation (“near gap”; s.d. = 22.5°) and the second flanker gap orientation was distributed 180° from the first flanker gap (“far gap”; s.d. = 22.5°). Because of the relatively narrow report error distributions even in the presence of a single flanker gap (e.g. Fig. 3A), we can with some confidence delineate which errors are associated with the near gap and which with the far gap. In Figure 4A, we show the raw report errors for the naïve observer. Report errors form two clusters: one cluster centred on the y-axis at approximately 0° and another at approximately ±180°. We arbitrarily defined reports with an absolute error greater than 90° as far-gap reports. These reports are shown above and below the top and bottom dashed lines, respectively, of Figure 4A. The proportion of far gap reports for each flanker size and each observer are shown in Figure 4B-D. For all observers, the proportion of far-gap flanker errors is greatest for the smallest flanker size, and gradually decreases as the flanker size increases.

**Figure 4.**
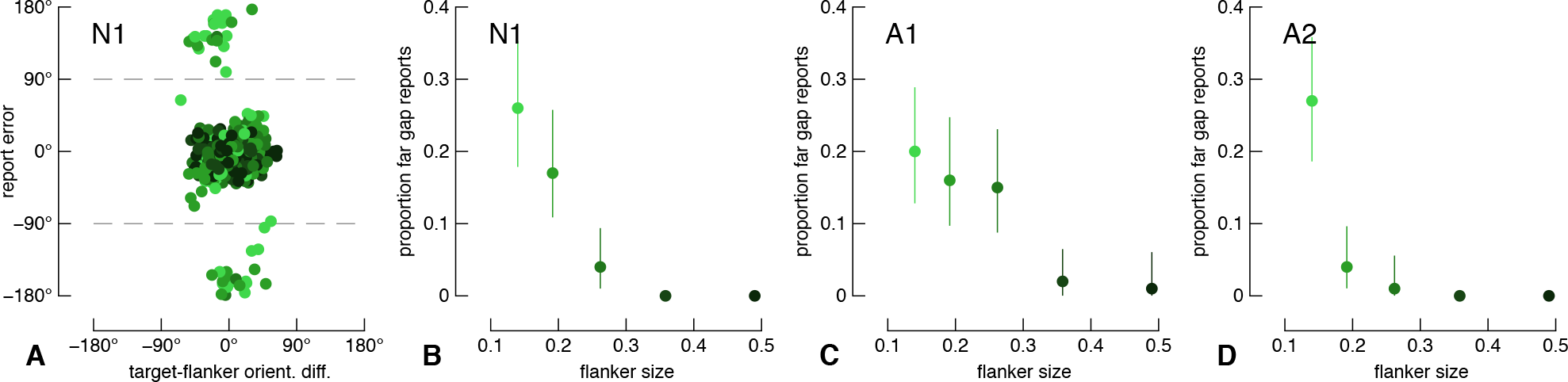
Report errors in the two-gap flanker condition. A) Raw report errors from the naïve participant. Report errors greater than 90° or less than -90° (dashed lines) were classified as trials in which observers misreported the far-flanker gap rather than the target or near-flanker gap. B-D) The proportion of such far gap reports as a function of flanker size for all observers. Errors bars are 95% confidence intervals.

We next divided the two-gap flanker condition data into two subsets for further analysis. First, we examined only those report errors within 90° of the target orientation. We performed the same analysis as we did for the one-gap flanker condition. Results are shown in Figure 5, and are highly similar to the results from the one-gap flanker condition in Figure 3. We also performed simple linear fits to the raw data, the results of which (misleadingly) favour an averaging model (see Appendix Fig. A2).

**Figure 5.**
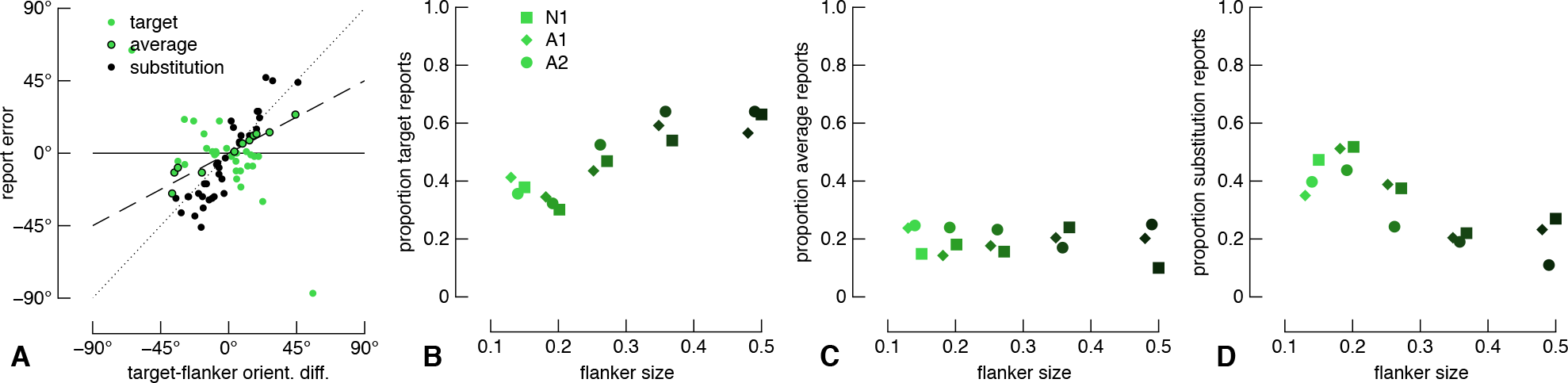
Results from the one-gap flanker condition. Data are shown as per Figure 3. A) Raw report errors from the naïve participant, focusing on only those errors within 90° of the target. B-D) Proportion of report type as a function of flanker size.

Second, we performed the same modelling on report errors that were greater than 90° from the target orientation. Due to the small number of observations in this analysis (Fig. 4), we pooled observers' data. We re-centered this subset of data by subtracting 180° from the orientation difference between the target gap and far flanker gap, as well as from the report error. The pooled errors corresponding to the far flanker gap are shown in Figure 6A. Because we re-centered these data, an error of 0° corresponds to a report of the target's polar opposite orientation, whereas data falling on the line of unity are reports following the far flanker gap orientation. As in the results above, with increasing flanker size, the proportion of target reports increases, the proportion of average reports is relatively stable, and the proportion of substitution reports decreases. In contrast to the results above, the proportion of target reports reached at the largest flanker size. However, there was only a single trial that had an error greater than 90° for the largest flanker condition, so this proportion necessarily had to be one or zero. Similarly, there were only two trials in the condition with the second largest flanker size, greatly restricting the possible proportions of report types.

**Figure 6.**
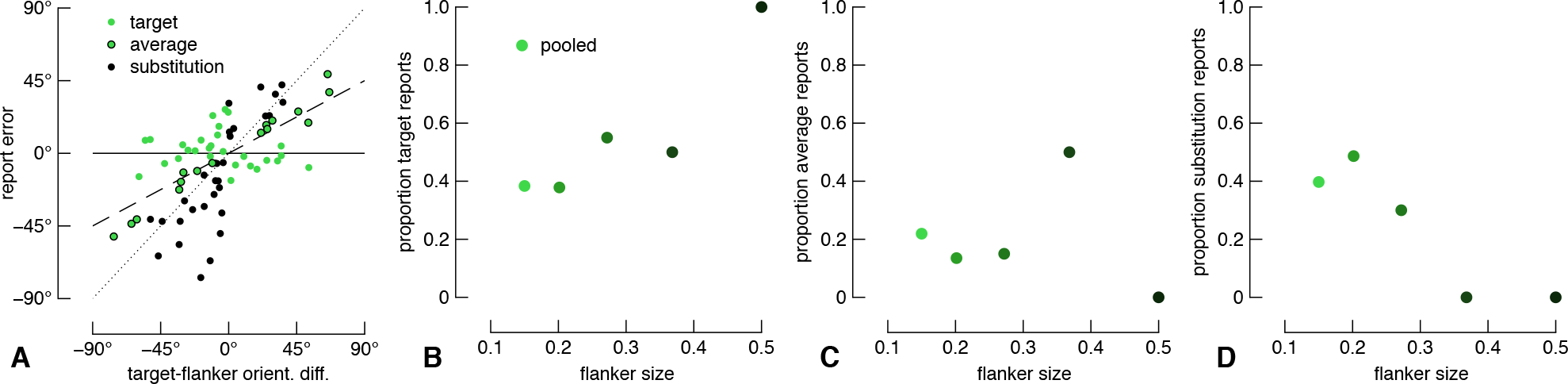
Results from the two-gap flanker condition for report errors greater than 90° from the target with re-centered data (see text). Data are shown as per Figure 3. A) Raw report errors pooled across observers. B-D) Proportion of report type as a function of flanker size.

## Discussion

We used a method of adjustment to quantify perceptual error in peripheral vision under novel crowded conditions. In all conditions, performance depended on the distance between the target and flanker, in line with the vast crowding literature ^4^. Based on the phenomenological responses and appearance of crowded stimuli, three general classes of mechanism have been advanced to account for crowding: 1) positional uncertainty ^30^, 2) feature averaging ^32^, and 3) source confusion ^33^. These models are not necessarily mutually exclusive and image processing based approaches have been advanced that incorporate elements of each of these mechanisms (refs 11 & 13), but it has been difficult to reconcile which best accounts for the data because of the use of different methodologies and stimuli across studies supporting each account. Furthermore, with image processing based models that produce foveal performance deficits with synthetic images that simulate peripheral vision, it is not clear which combinations of these underlying processes accounts for perceptual performance ^11,51,52^. The experimental design and complementary modelling employed here provides a novel way to classify the frequency of each error type with the same stimuli and observers. Our results reveal that all error types characterise crowding with Landolts, but their proportions vary with the distance between the target and flanking stimuli.

In the no-gap flanker condition, the crowding we observed at relatively small target-flanker separations is likely caused exclusively by an increase in orientation uncertainty (for example, ref ^30^). In this condition, there are no flank features for averaging or substitution to occur, yet, as shown in Figure 2, we found a reliable increase in the circular standard deviation of perceptual errors with decreasing target-flanker separation, see also ^37^. This result is unlikely due to a form of classical masking, such as meta-contrast masking, because perceptual reports were not randomly distributed in the presence of close flankers ^53^. It is also difficult to account for these errors with an attentional account of crowding, in which observers' attentional resolution is too coarse to individuate the target gap ^5^: even in the smallest flanker condition, observers' reports cluster around the actual target position, indicating that they could indeed attend to the target gap, albeit with greater perceptual error that is directly attributable to an increase in orientation variance (Fig. 2). We suggest that this orientation noise can be attributed to the solid flanking ring increasing the bandwidth of a population code that encodes the target orientation (see below and ref ^37^).

The results of the one-gap and two-gap flanker conditions also support recent proposals that crowding may best be accounted for by a population code. Rather than conforming neatly to a single report error type, we found errors could be accurate, follow the flanker gap, or some average of the two. These data are thus difficult to reconcile with simple averaging or substitution models. Van den Berg et al ^47^ showed that many hallmarks of crowding can be explained by a biologically inspired model that simulates the responses of populations of neurons tuned to orientation within a fixed region of space (ie. a receptive field). Crowded stimuli create systematic shifts in the population code, so that when the population code is decoded, the decoded signal is prone to error. We further showed that an idealised population code can, for a given crowded stimulus, produce accurate, averaged, or substituted report errors in a probabilistic fashion ^37^. We thus argued that there is no averaging or substitution mechanism per se, but instead that perceptual reports are drawn from the population response to the stimuli. Such a process is distinct from any single phenomenon such as source confusion, averaging or substitution but instead results from the broad spatial bandwidth of early stage filters.

Critically, the results from the present study show that averaging of features is not compulsory, in contrast to previous work ^32^. On a number of trials in which a flanker gap is present, observers can recover the target orientation with a precision similar to that observed with no flanker gap (Fig. 3). With only a single report, it is impossible to know if, on a single trial, an observer perceived both the target orientation and the flanker orientation, their average, or if the frequencies of these percepts varied across trials. We previously asked participants to report both the target and flanker orientations in an experiment similar to the present report, and found that participants were generally capable of reporting both elements, though they often reversed feature positions ^37^. Taken together, these data reveal that perceptual reports in clutter are probabilistic, but relatively fine detail can be recovered.

It is clear from our data that observers often report an orientation closer to the flanker orientation instead of the target orientation (Fig. 4). However, our findings suggest the proportion of substitution-type errors is substantially greater than the proportion of substitution errors that occur when whole letter stimuli are used ^39^. It is likely that this discrepancy can be explained by the task differences. Letter report paradigms limit the response range and so the observer is forced to select the most similar letter, even if the perceived stimulus does not match any of the possible responses. Our response method does not suffer from this limitation.

Our results thus provide new evidence revealing that the component features of visual objects can be individuated even far in the periphery, although their relative positions and orientations may appear noisy and confusable across trials. The level of detail made available by the visual system has been heavily debated both within the crowding literature and more generally. Our data suggest one possible reason for this apparent conflict in the literature. Note that our analyses suggest all observers have very high rates of substitution-type reports under crowded conditions: combining the proportion of far-flanker gap substitutions (Fig. 4) and proportion of near-gap errors (Fig. 5), observers made approximately 50% - 70% substitution-type errors in the most crowded conditions. Such performance may be misinterpreted in alternative forced choice experiments (AFC), a more common psychophysical paradigm in which an observer is forced to categorise a target as being one of usually two to four alternatives. In these experiments, such a high proportion of substitution-type reports could render performance at or close to chance, leaving it unclear if a participant was randomly guessing, reporting an average stimulus or reporting what they thought was the target on some trials and the flanker on other trials. This is especially true in experiments with simple stimuli such as lines or Gabors, though it is less problematic with letter stimuli e.g. ^34,39^.

In conclusion, our findings show that relatively fine featural detail is not necessarily lost during early visual processing, but the precision of each perceptual report is corrupted. The aggregate of errors made when viewing crowded displays cannot be characterised as simply being accurate, averaged or substituted. The variety of report error types that occur within the same paradigm, as demonstrated here, provides a continued challenge to models of visual crowding.

## Author Contributions

Both authors designed the experiments and analyses. WJH collected the data and wrote the manuscript. PJB and WJH analysed the data, and PJB revised the manuscript.

## Acknowledgements

This work was supported by NIH grant R01EY021553 (P.J.B.) and a National Health and Medical Research Council of Australia CJ Martin Fellowship (APP1091257; W.J.H.).

The authors declare no competing nancial interests.

## APPENDIX A Supplementary Analyses

Prior to the mixed-model analyses for the one-gap and two-gap flanker conditions presented in the Results section, we performed the following linear analyses.

### One-gap flanker condition

Shown in Figure A1A are the naïve observer's report errors and linear fits as a function of target-flanker orientation difference. Note that a linear fit with a slope of 1 would indicate the observer's reports followed the flanker gap closely, a slope of 0 would indicate the observer's reports were not influenced by the flanker, and a slope of 0.5 conforms to the average of target and flanker orientations. In Figure A1B-D we plot the slope parameter for all conditions from all observers for the one-gap flanker condition. Slope parameters were close to 0.5 for the two smallest flanker sizes, and generally decreased as the flanker size increased. For all observers, the slopes were approximately 0 for the largest flanker condition. However, these analyses do not describe the data in full (see Results).

**Figure A1.**
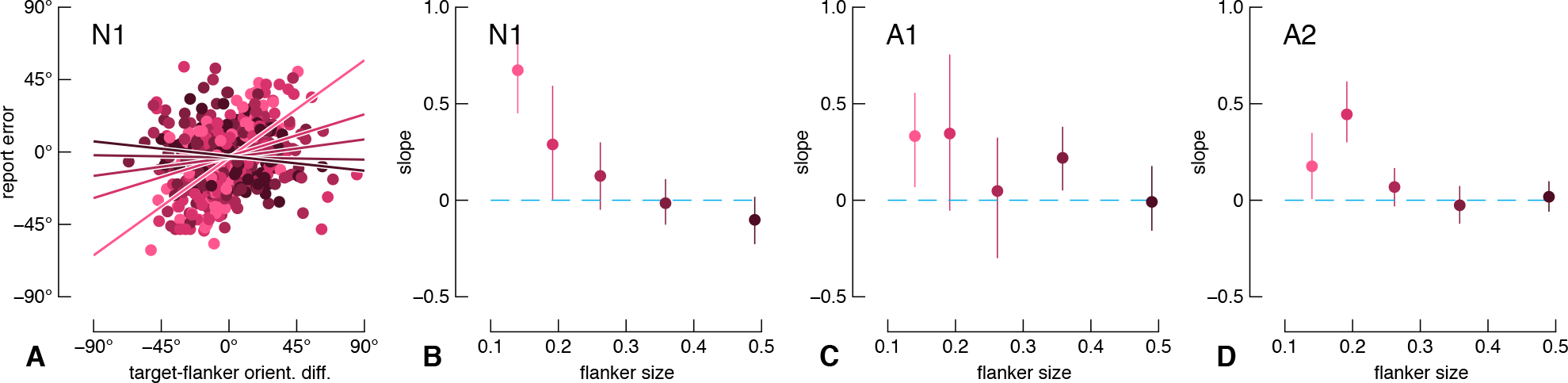
Results from linear fits in the one-gap flanker condition. A) Example raw data from the naïve participant. Lines show linear fits for each target-flanker separation, with colour shading indicating conditions as in Fig. 2A. To improve visibility, we have truncated the x-axis and y-axis. B-D) Slope parameters as a function of flanker size for all observers. Errors bars are 95% confidence intervals.

### Two-gap flanker condition (near-gap)

**Figure A2.**
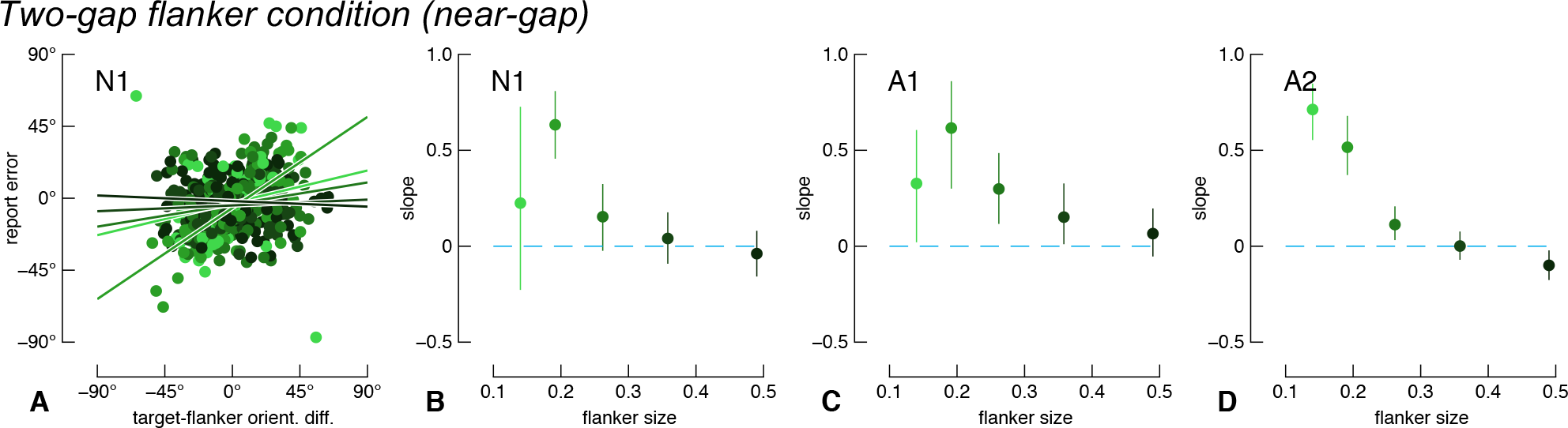
Results from the two-gap flanker condition for report errors less than 90° from the target. A) Raw data from the naïve participant. The x-axis indicates the orientation difference between the target gap and the nearest flanker gap. Data presented as in Fig. A1A. B-D) Slope parameters as a function of flanker size for all observers. Errors bars are 95% confidence intervals.

## Footnotes

1 Note that our target is a modified version of a Landolt C. A true Landolt C has a gap section formed by parallel lines, whereas the gap section in our stimulus is formed by non-parallel lines.

2 http://uk.mathworks.com/matlabcentral/fileexchange/10676-circular-statistics-toolbox–directional-statistics-

